# Formation of wall-less cells in *Kitasatospora viridifaciens* requires cytoskeletal protein FilP in oxygen-limiting conditions

**DOI:** 10.1101/2020.05.26.116947

**Authors:** Eveline Ultee, Ariane Briegel, Dennis Claessen

## Abstract

The cell wall is considered an essential component for bacterial survival, providing structural support and protection from environmental insults. Under normal growth conditions, filamentous actinobacteria insert new cell wall material at the hyphal tips regulated by the coordinated activity of cytoskeletal proteins and cell wall biosynthetic enzymes. Despite the importance of the cell wall, some filamentous actinobacteria can produce wall-deficient S-cells upon prolonged exposure to hyperosmotic stress. Here we performed cryo-electron tomography and live cell imaging to further characterize S-cell extrusion in *Kitasatospora viridifaciens*. We show that exposure to hyperosmotic stress leads to DNA compaction, membrane and S-cell extrusion and thinning of the cell wall at hyphal tips. Additionally, we find that the extrusion of S-cells is abolished in a cytoskeletal mutant strain that lacks the intermediate filament-like protein FilP. Furthermore, micro-aerobic culturing promotes the formation of S-cells in the wild-type, but the limited oxygen still impedes S-cell formation in the Δ*filP* mutant. These results demonstrate that S-cell formation is stimulated by oxygen-limiting conditions and dependent on the presence of an intact cytoskeleton.

## INTRODUCTION

Almost all bacteria are enveloped by a cell wall that provides protection against challenging or hostile environments. The cell wall also prevents cells from bursting by withstanding the high internal turgor pressure. Furthermore, it is a major determinant of bacterial cell shape. A main constituent of the cell wall is peptidoglycan (PG), which consists of cross-linked glycan strands that create in essence a single gigantic molecule called the sacculus ^1^. The biosynthesis of PG is conserved in bacteria and is therefore a prominent target for many clinically used antibiotics. Cell wall precursors are synthesized in the cytosol and are subsequently transferred to the exterior of the cell. Here, designated synthases incorporate these monomers into the pre-existing cell wall ^2–4^.

The site of incorporation of new cell wall material is species-specific and occurs over the entire surface of the lateral wall in many rod-shaped bacteria, while incorporation is confined to apical sites in filamentous actinobacteria. Tip growth in filamentous actinobacteria is orchestrated by the TIPOC (<u>tip</u> <u>o</u>rganizing <u>c</u>enter) multiprotein complex ^5–8^. The best-studied component of this complex is the essential protein DivIVA, which drives apical growth by recruiting the cell wall synthesis machinery and marks the location of new hyphal branch sites ^5,7,9^. In order for the cell wall to grow, the existing PG needs to be modified to allow new monomers to be incorporated and cross-linked into the sacculus, making the hyphal tip a putative vulnerable structure. Previously, the intermediate filament-like protein FilP was proposed to reinforce the tip intracellularly ^10^. FilP localizes to active growing hyphal tips, where it forms cytoskeletal structures that interact with DivIVA ^8^.

Although the cell wall is considered an essential structure in bacteria, many species can shed their cell wall to overcome PG targeting threats, such as antibiotics and the mammalian immune system ^11–14^. In laboratory conditions, the transition from a walled state to the cell wall-deficient (CWD) state is typically induced by exposing bacteria to PG synthesis-targeting antibiotics and/or lytic enzymes ^15,16^. Our lab has previously shown that several filamentous actinobacteria have a natural ability to form CWD cells without the help of PG synthesis-targeting compounds ^17^. These CWD cells, termed S-cells for <u>s</u>tress-induced <u>cells</u>, are extruded from hyphal tips in hyperosmotic environments following an arrest in tip growth. S-cells are unable to proliferate without their cell wall but can sustain in their CWD state for prolonged periods of time and thus likely survive stressful conditions.

Occasionally, tip growth is reinitiated from branches that appear subapically from the stalled tip after extrusion of the S-cells. This observation implies that the filament from which the S-cells emerge is still viable. How S-cells are extruded and how this process is regulated at the molecular level is poorly understood.

In this study, we combine genetics with fluorescence time-lapse microscopy and cryo-electron tomography (cryo-ET) to characterize the morphological and structural changes associated with S-cell formation. Our data reveal that oxygen limitation triggers S-cell formation in the wild-type strain in a FilP-dependent manner. These results suggest that S-cell extrusion is a controlled physiological adaptation to stress and depends on cytoskeletal elements involved in polar growth.

## RESULTS

### Membrane and DNA organization during S-cell extrusion

We previously showed that prolonged exposure to hyperosmotic stress causes an increase in branching frequency, membrane synthesis and DNA condensation in *K. viridifaciens* ^17^. To characterize these changes in more detail, we performed time-lapse microscopy in combination with fluorescent dyes that bind to nucleic acids and lipids (SYTO9 and FM5-95, respectively). Time-lapse imaging of growing filaments indeed revealed condensed DNA and an excess of membrane in high osmotic conditions (Supplementary Movies 1A, B). Strikingly, excess membrane was frequently extruded from the hyphal tips of both leading tips and emerging branches (Supplementary Movie 1A, Fig. 1, arrowheads). Regrowth of the hyphal tip is associated with strong turns or bends, which could indicate a local rearrangement of the TIPOC leading to a new growth direction (Supplementary Movie 1B). In some cases, the membrane that blebs off from the hyphal tip enlarges and forms large vesicles with a diameter of 4-5 μm (Fig. 1, asterisk in 6h00 panel). Some of these vesicles emit green fluorescence, indicating the presence of SYTO9-stained nucleic acids and therefore we consider them S-cells. Subsequently, extruded smaller vesicles at the same tip are typically smaller and often lack nucleic acids (Fig. 1, arrows in 7h00 panel). The hyphae still possess DNA after extruding S-cells. This could indicate that either DNA replication is ongoing, or that the nucleoid is changing its organization and morphology upon exposure to stress.

**Figure 1.**
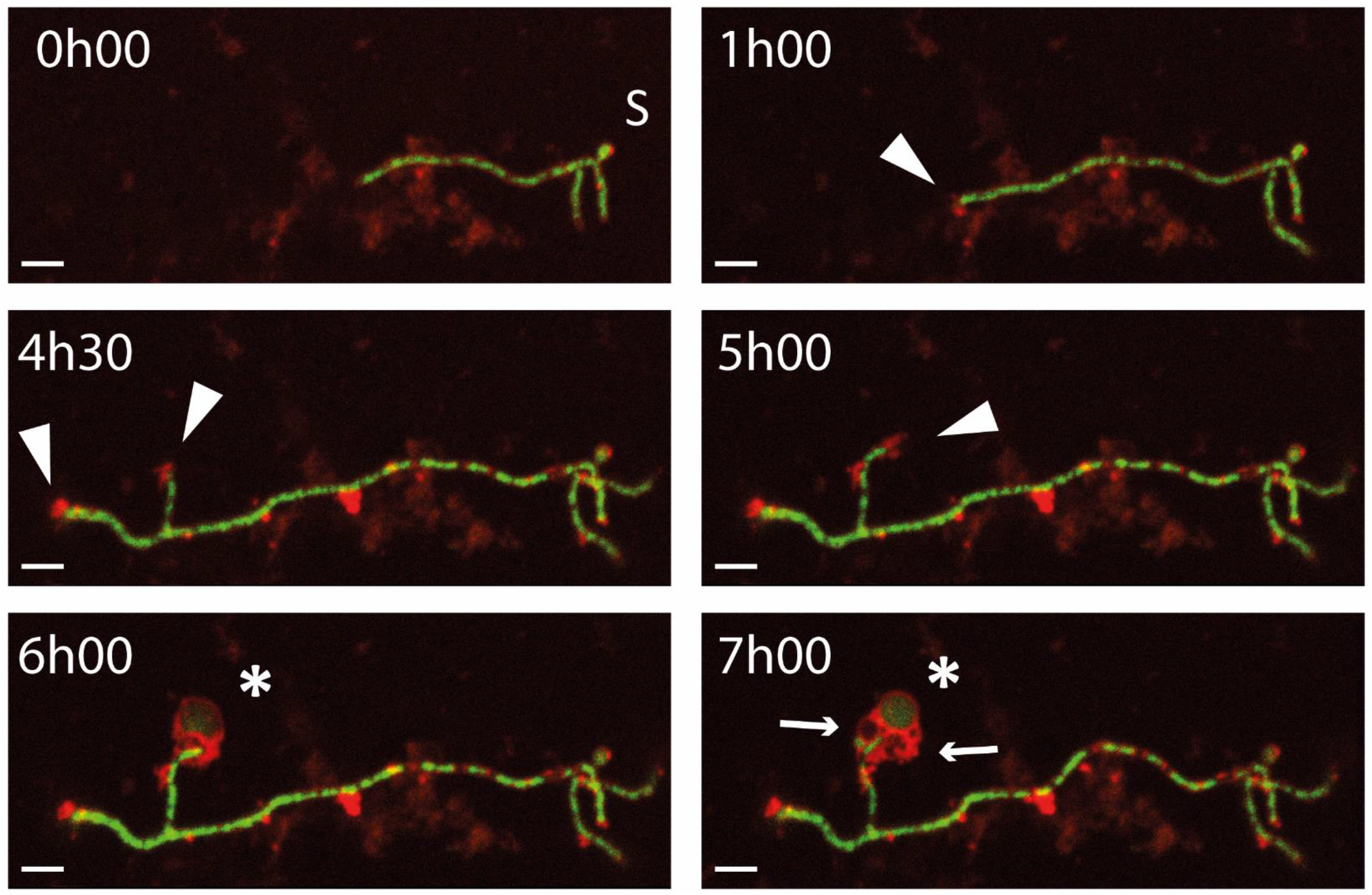
Extrusion of S-cells from *K. viridifaciens* germlings under high osmotic stress. Germinated *K. viridifaciens* spores were fluorescently labelled with SYTO9 (nucleic acids) and FM5-95 (lipids) and were grown under high osmotic conditions. Micrographs were taken every 30 minutes (see Supplementary Movie 1) of which a selection of micrographs are shown. The ‘S’ indicates the spore, the arrowheads highlight the blebbing of membrane, the asterisk indicates the S-cell and the arrows show the formation of membrane vesicles. Scale bar: 5 μm

To quantify these observations, we categorized and quantified the various events associated with the formation and release of S-cells. In order of appearance, these stress response-related events are: (B) membrane extrusion without the formation of obvious membrane vesicles, (C) formation of empty membrane vesicles, (D) formation of DNA-containing S-cells, and (E) the presence of both S-cells and empty membrane vesicles (Fig. 2). Quantitative analysis shows that after 18 hours of exposure to osmotic stress, on average 71% of the hyphal tips showed one of the abovementioned S-cell-related stress responses (responses B-D), while 29% of the tips appeared not affected (A). 15% of all imaged and analyzed tips showed excessive membrane blebbing, while DNA-containing S-cells were observed surrounding 44% of the hyphal tips (Fig. 2, Supplementary Table 1). The majority of tips surrounded by S-cells also showed empty vesicles and/or excessive membrane blebbing. Of three experimental replicas, the number of non-responding and blebbing hyphal tips shows relatively little variance, whereas the occurrence of empty and/or filled vesicles differs per replica (Supplementary Table 1).

**Figure 2.**
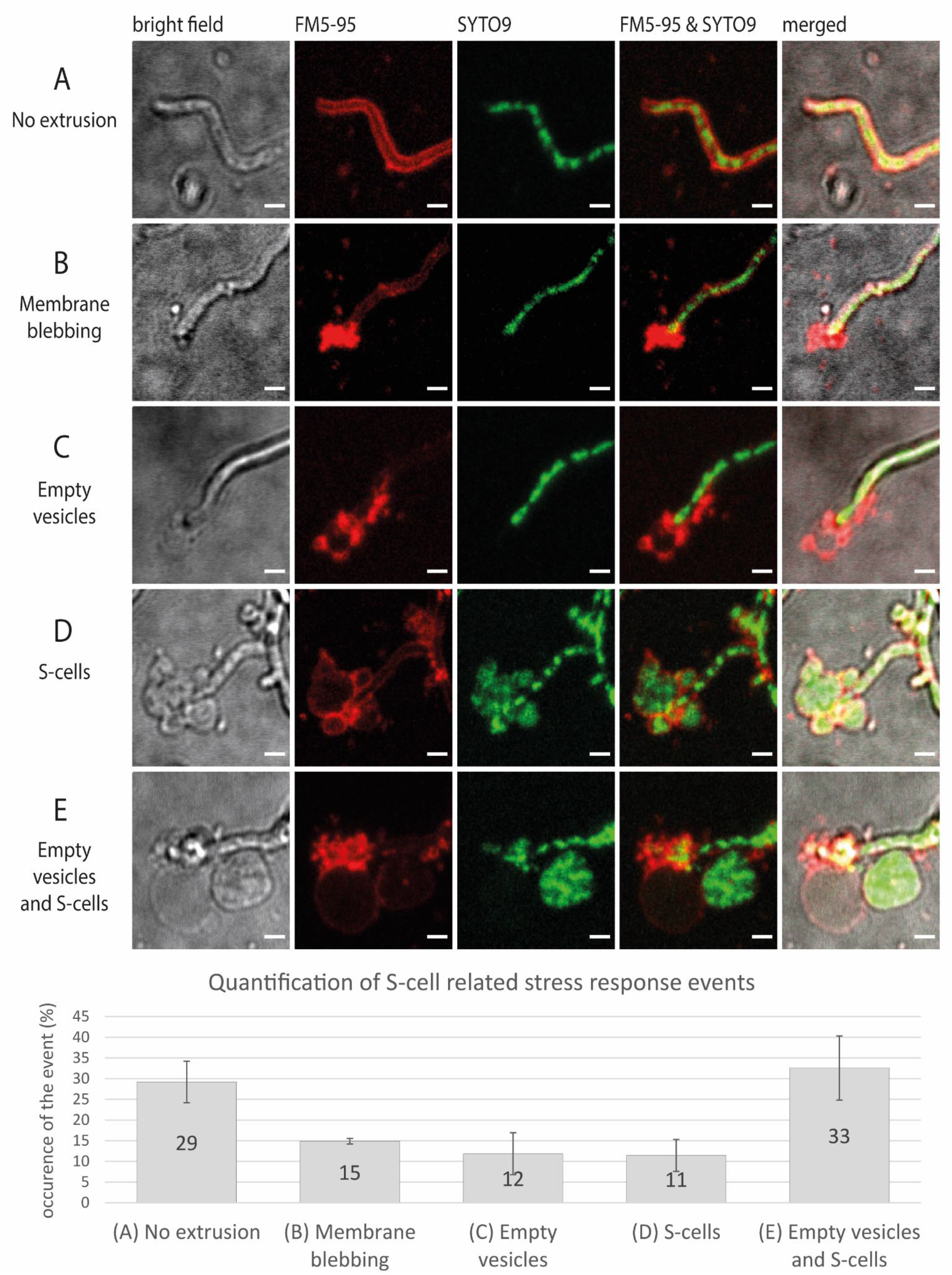
Quantification of S-cell-related stress response events at the hyphal tip. Germinated *K. viridifaciens* spores were fluorescently labelled with SYTO9 (nucleic acids) and FM5-95 (lipids) and were grown under high osmotic conditions. After 18 hours of growth, the hyphal tips were imaged (representative micrographs shown), examined and categorized into (A) no visible S-cell-related stress responses, (B) membrane blebbing tips, (C) the presence of empty vesicles, (D) the presence of S-cells or (E) the presence of empty vesicles and S-cells. The average of three experimental replicas is shown in the graph (data in Supplementary Table 1). Scale bars represent 2 μm.

### Structural changes in the cell envelope accompany S-cell extrusion

In order to observe the structural changes in the hyphal tip during S-cell release, cryo-transmission electron microscopy (cryo-TEM) was used (Fig. 3A, B). Interestingly, the cell envelope thickness, measured from the cytoplasmic membrane to the outer edge of the cell wall, was considerably thinner when *K. viridifaciens* was grown in medium supplemented with 20% sucrose: 31 nm (±1,14; n=15), compared to an average of 43 nm (±1,05; n=15) when grown in the same medium without sucrose (Supplementary Table 2).

**Figure 3.**
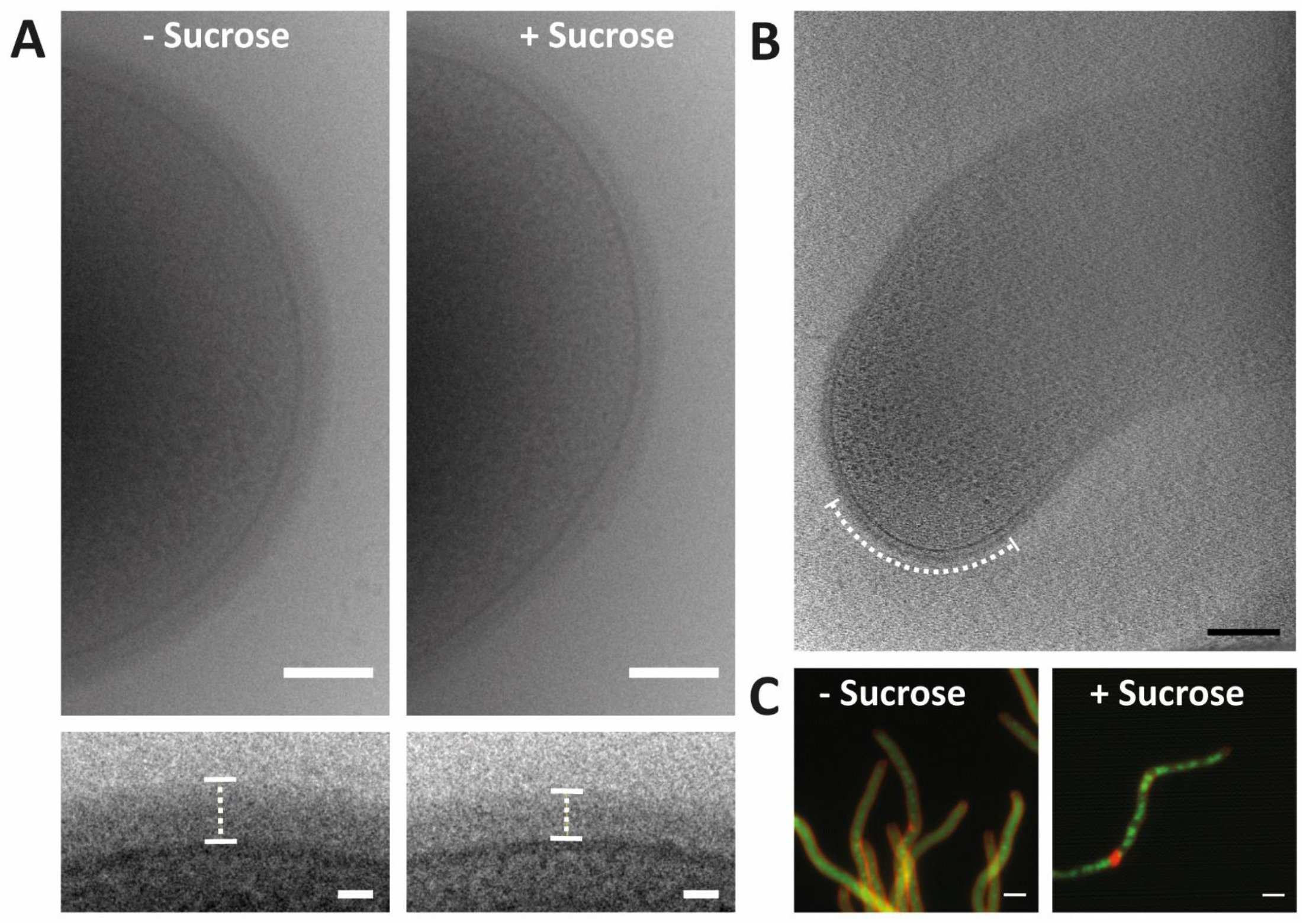
Structural changes in *K. viridifaciens* hyphae upon exposure to osmotic stress. Cryo-TEM micrographs of the tip of *K. viridifaciens* hyphae grown with or without 20% sucrose show a difference in cell wall thickness, highlighted in the insets (A). Cryo-ET micrograph of a 17-nm thick slice of *K. virdifaciens* grown with osmotic stress shows a ribosome excluded zone at the terminus of the tip (B). Fluorescently-stained *K. virdififaciens* hyphae (lipids: FM5-95, red) show absence of nucleic acids at the tip and condensation of the DNA (SYTO9, green) upon exposure to osmotic stress (C). Scale bars represent 100 nm (upper panels A), 20 nm (lower panels A), 200 nm (B) or 5 μm (C).

Osmotically-stressed *K. viridifaciens* cells were further investigated with cryo-electron tomography (cryo-ET), allowing three-dimensional visualization of the filaments and S-cells at high-resolution. Despite the sub-optimal thickness of the hyphal cells for penetration of the electrons (>500 nm) and sub-optimal buffer (20% sucrose, resulting in low contrast) we were able to detect structural elements. Ribosomes can be distinguished inside the hyphal cells but are absent in the extreme end of the tip (Fig. 3B). In fact, roughly 200-400 nm of the tip’s apex does not appear to contain ribosomes or DNA. Fluorescent labeling of the chromosome in both stressed and non-stressed filaments confirm the absence of the chromosome at the extreme end of the tip (Fig. 3C). We also analyzed extruded S-cells and membrane vesicles with cryo-ET. In line with the light microscopy data, the released S-cells ranged in size and content. While some vesicles contain ribosomes, DNA and/or inner vesicles (Fig. 4A, B), others appear empty (Fig. 4C). Notably, all imaged S-cells lacked a visible cell wall, and were comparable to cells of a penicillin-induced L-form strain of *K. viridifaciens* ^17^ (Fig. 4D).

**Figure 4.**
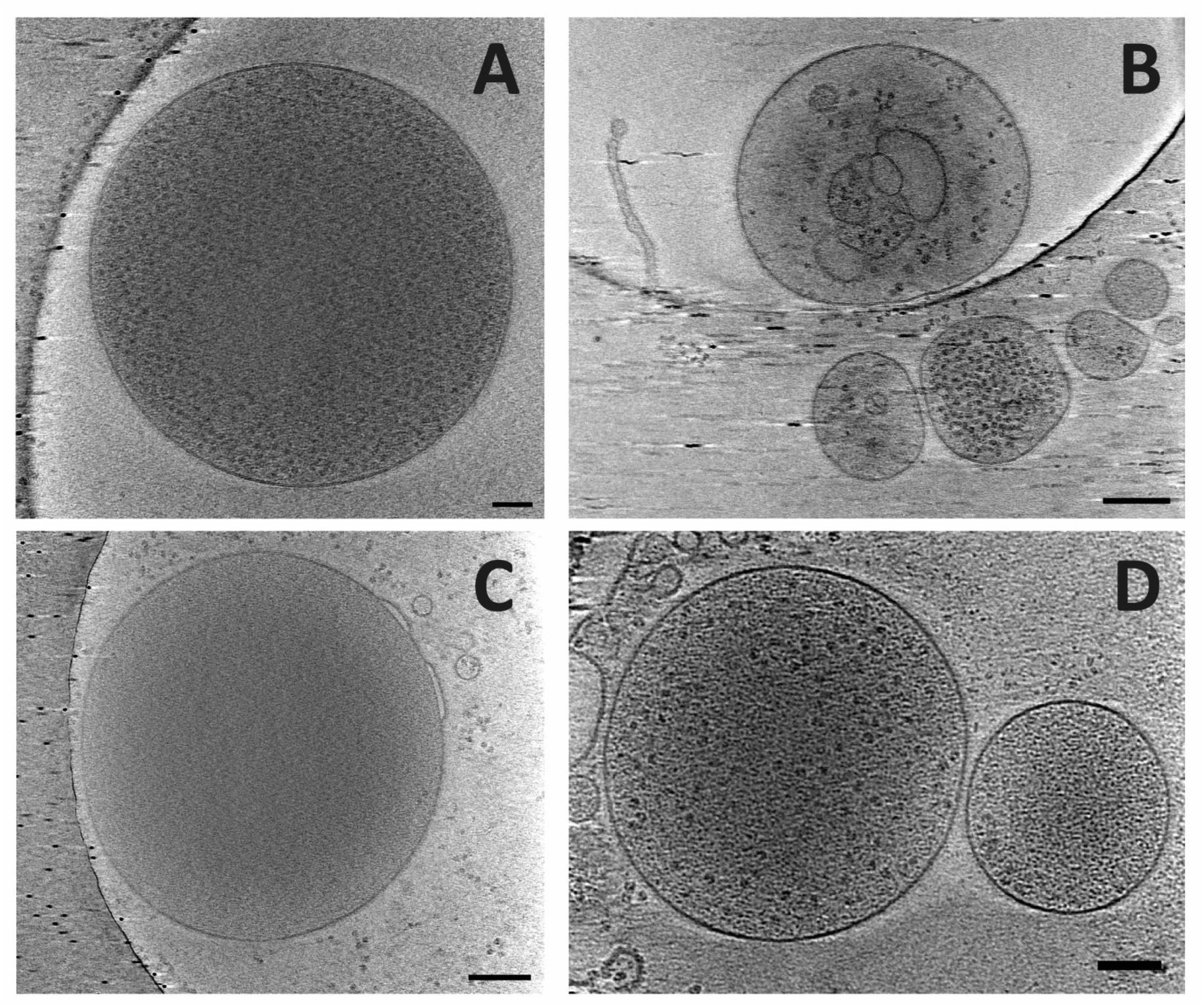
Heterogeneity among *K. viridifaciens* S-cells. Cryo-ET micrographs show S-cells formed by young hyphae of *K. viridifaciens* after 6 hours of exposure to high sucrose levels (A-C). The S-cells structurally resemble L-forms (D). Scale bars represent 100 nm (panels A and D) or 200 nm (panels B and C).

### FilP is required for S-cell formation

Our data shows that the hyphal tip undergoes a visible structural transition prior to S-cell release. Since the cytoskeletal protein FilP localizes to actively growing hyphal tips and is linked to the sturdiness of the tip in *Streptomyces* ^10^, we reasoned that it could play a significant role in the release of S-cells from osmotically stressed hyphae. We therefore created a knock-out mutant strain of the *filP* homologue in *K. viridifaciens* (BOQ63_RS19300 ^18^, Supplementary Fig. 1). First, the morphology of the *K. virdifaciens ΔfilP* strain was studied using time-lapse microscopy while grown underneath an agar pad, composed of MYM (no osmotic stress) or LPMA (high osmotic stress) medium (Supplementary Movies 2-5). The wild-type and Δ*filP* strains both grow equally well in the absence of osmotic stress (Supplementary Fig. 2, Supplementary Movies 2, 3). In contrast, when the strains were exposed to hyperosmotic stress, the wild-type was able to extrude S-cells (Fig. 5A, Supplementary Movie 4) while the Δ*filP* strain was not. Instead, the mycelium of this mutant started to form numerous small side branches and adopted a highly compact morphology (Fig. 5A, Supplementary Movie 5). Further investigation using fluorescent dyes showed that not only the formation of S-cells was impaired in the Δ*filP* mutant, but also the extrusion of excess membrane (Fig. 5B). This is in stark contrast to the wild-type strain, which produce an abundance of S-cells, vesicles and extruded membrane (Fig. 2, 5B, Supplementary Table 1). Interestingly, the morphology of the Δ*filP* mutant was comparable to the wild-type when the strain was grown on LPMA plates without being covered by an agar pad (Supplementary Fig. 3). Under these conditions, the Δ*filP* mutant was able to generate S-cells, but again failed to do so when an agar pad was placed on top of the strain (Supplementary Fig. 4). The Δ*filP* mutant was also able to extrude S-cells when grown under slow shaking conditions in liquid LPB medium, which, like LPMA, contains 20% sucrose (Supplementary Fig. 5). Altogether, these data identify FilP as an important protein involved in S-cell extrusion. It furthermore demonstrates that the inability of the Δ*filP* mutant to generate S-cells is not exclusively related to hyperosmotic stress.

**Figure 5.**
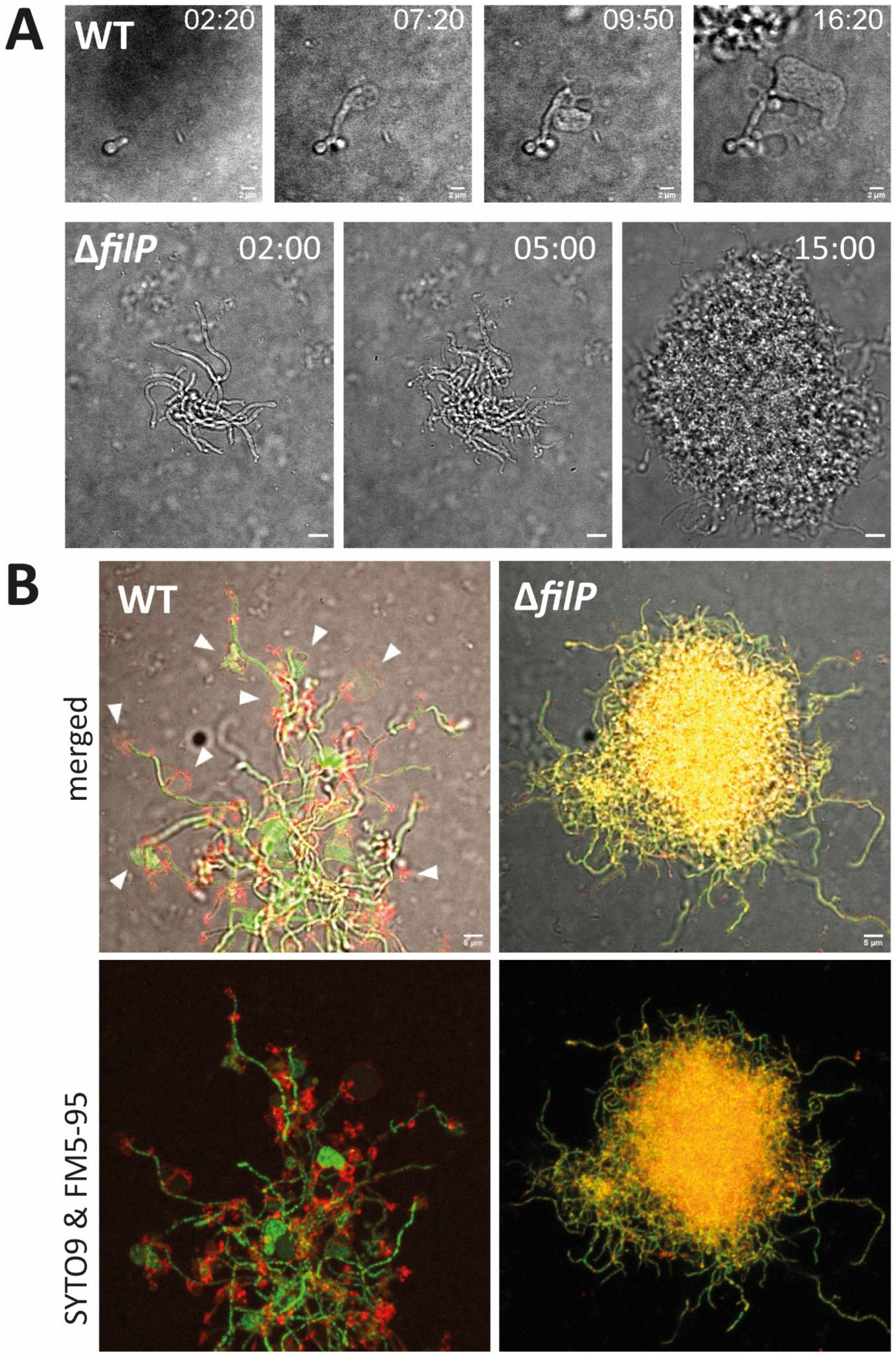
FilP is required for S-cell extrusion in *K. viridifaciens*. (A) Time-lapse microscopy stills of *K. viridifaciens* germlings under LPMA agar pads. Wild-type germlings (top panels) show S-cell formation, whereas those of the Δ*filP* strain do not form S-cells (bottom panels). Instead, the Δ*filP* strain hyper-branches leading to the formation of dense mycelial pellets (see Supplementary Movie 5). (B) Micrographs of germinated *K. viridifaciens* wild-type and Δ*filP* spores that were fluorescently labelled with SYTO9 (nucleic acids, green) and FM5-95 (lipids, red) and grown underneath an LPMA agar pad. Images were taken after 18 hours of growth. Arrows highlight the formation of S-cells. Time is indicated in the hours:minutes format and scale bars represent 2 μm (A) or 5 μm (B).

### FilP is required for S-cell formation in oxygen-limiting conditions

Our results indicated that FilP is important for the formation of S-cells in conditions when the strain was grown in a confined space in between two layers of agar. We reasoned that this setup could potentially reduce the levels of oxygen that is accessible to the cells. To further study the effect of oxygen limitation on S-cell formation, the *K. viridifaciens* wild-type and Δ*filP* strains were grown on solid LPMA and MYM medium in micro-aerobic vessels. Both the wild-type and Δ*filP* strains grew slowly in micro-aerobic conditions but were able to germinate and establish a mycelium after 14 days of growth (Fig. 6, Supplementary Fig. 6). On MYM medium, the wild-type strain formed a regular-looking mycelium consisting of branching hyphae. Although, some occasions of apical branching or tip splitting could be observed (Fig. 6, asterisks) which indicates a distortion in the TIPOC functioning ^6^. In contrast, the Δ*filP* mutant did not show obvious tip splitting, and formed short and apparently thick mycelial fragments that appear to have a lower branching frequency (Fig. 6). Importantly, when cultivated micro-aerobically and exposed to osmotic stress, the wild-type strain formed an excessive number of S-cells and vesicles that varied in size. Moreover, wild-type hyphal tips often showed signs of bulging reminiscent of sporulation (inlays in Fig. 6). Bulging of hypha was also observed in the Δ*filP* strain, but the mutant strain was unable to form S-cells under these micro-aerobic conditions. Altogether, these data indicate that in oxygen-limiting conditions, FilP is required for the extrusion of S-cells, as well as for the establishment of a normal mycelial architecture in the absence of osmotic stress.

**Figure 6.**
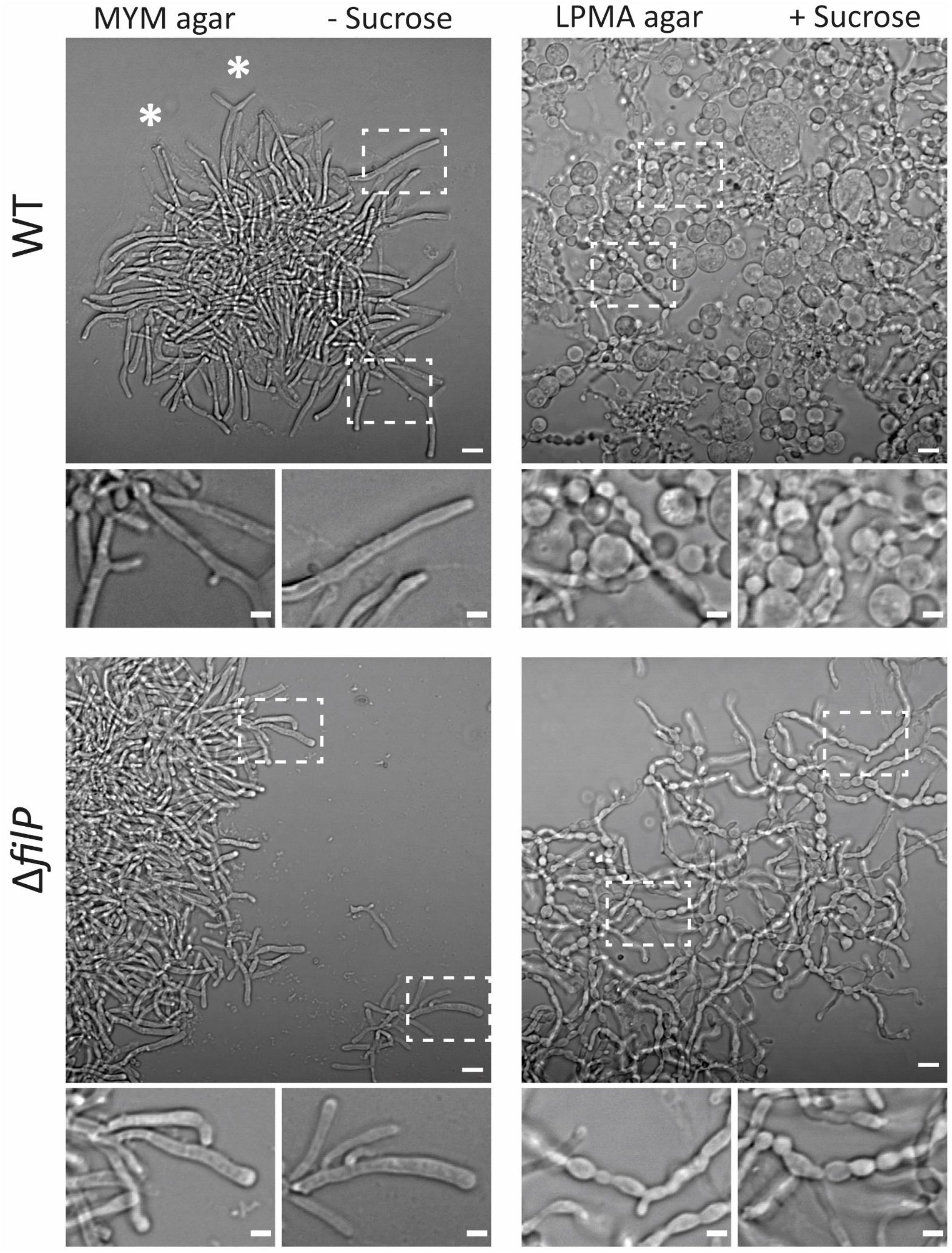
Morphology and S-cell extrusion of *K. viridifaciens* in micro-aerobic conditions. Spores of the *K. viridifaciens* wild-type and Δ*filP* strains were cultured micro-aerobically on MYM (no osmotic stress) and LPMA (high osmotic stress) solid medium. Filaments of the wild-type strain show tip-splitting on MYM medium (asterisks). S-cells and vesicles are present in the wild-type culture on LPMA but are absent in the cultures of the Δ*filP* mutant. The inlays show bulging hyphae in both LPMA-grown cultures. Scale bars represent 5 μm or 2 μm (inlays).

## DISCUSSION

We previously showed that cell wall-deficient S-cells are formed by filamentous actinomycetes upon exposure to hyperosmotic stress, but the underlying regulation of this process was unknown. In this study, we reveal the detailed structural changes that accompany S-cell formation and release for the first time. Furthermore, our results show that the absence of cytoskeletal protein FilP distorts hyphal morphology and blocks S-cell formation. S-cell formation appears to increase in the wild-type strain when grown in micro-aerobic conditions but remains absent in the strain lacking FilP. These results highlight the importance of FilP for the extrusion of S-cells and identifies FilP as the first protein to be involved in the regulation of S-cell formation in *K. viridifaciens*. The increased formation of S-cells in oxygen-limiting conditions further suggests that S-cells may serve an important function in oxygen-deprived environments.

Filamentous microorganisms incorporate new wall material at the hyphal tips. In such organisms, turgor pressure is important for growth and believed to physically stretch the wall to allow incorporation of cell wall precursors ^19–22^. Hyperosmotic environments lower the turgor pressure and could therefore reduce the speed of apical wall synthesis. Additionally, this could also induce plasmolysis through the separation of the cytoplasmic membrane from the bacterial cell wall ^23–25^. Both events could contribute to S-cell formation and their release from the tip. Potentially, if cell wall synthesis but not membrane synthesis is affected by changes in turgor pressure, this could lead to a disbalance and relative overproduction of membrane.

Our cryoET data shows that the cell wall is thinner when *K. viridifaciens* is exposed to hyperosmotic stress. Indeed, this would be expected given that osmotic upshifts lead to a growth arrest at the leading hyphal tip ^26^. As a consequence, less nascent PG is inserted and might result in a thinner and structurally weaker cell wall. As nascent PG is exclusively incorporated at the hyphal tip, this forms the weakest spot of the hyphal cell wall ^27^ and explains why S-cells are extruded at the tip. The extrusion furthermore appears to depend on FilP, which is part of a stress-bearing cytoskeleton. We hypothesize that the necessary turgor pressure for extrusion of S-cells becomes too low in the *filP* mutant in hyperosmotic conditions.

### A role for oxygen deprivation in S-cell extrusion

Aerobic bacteria use oxygen as an electron acceptor in the electron transfer chain (ETC), which causes an increase in the levels of reactive oxygen species (ROS). Increased ROS levels appear to be an indirect result of blocking PG synthesis, leading to an excess of sugars metabolized via the TCA cycle and consequently more ROS generated as by-product in the ETC. Notably, L-form bacteria were found to be sensitive to oxidative damage caused by high ROS levels ^28^. Counteracting these high amounts of cytotoxic ROS compounds, as by redirecting carbon metabolism, allows the CWD cells to become robust and proliferating L-forms ^28,29^. When grown anaerobically, oxygen is not available as electron acceptor in the ETC chain and therefore ROS levels are lower. We here observed that S-cells appear more abundant when *K. viridifaciens* is grown in micro-aerobic conditions. Whether the reduced oxygen levels facilitate the extrusion process or even lead to proliferation of S-cells is unknown.

Furthermore, the absence of FilP has a dramatic effect on the ability to form S-cells under oxygen stress. To our knowledge, no direct link between the apical growth machinery and oxygen stress has been described before. The deviant tip morphology of the wild-type strain on growth medium with limited oxygen supports the hypothesis that oxygen levels influence the TIPOC. However, limited knowledge is available on micro-aerobic growth in filamentous actinomycetes, which may in fact be a condition that these bacteria frequently encounter in the heterogeneous soil environment.

As soil-dwelling bacteria, filamentous actinomycetes are often found in association with eukaryotic hosts, including plants ^30^. The root systems of plants are highly dynamic in their oxygen availability and can, depending on their abiotic influences, transition to an anoxic state ^31^. Interestingly, CWD cells or L-forms have been observed inside plant tissues. It has been hypothesized that the transition from intercellular to intracellular invasion leads to a loss of the cell wall in root tissue ^32,33^. Strawberry plants inoculated with *B. subtilis* L-forms showed that these bacterial cells can travel from the injection spot to up to 42 cm through the plant ^34^, a task that would be difficult to achieve by non-motile, filamentous bacteria such as *K. viridifaciens* or *Streptomyces* species. Here we show that micro-aerobic conditions promote S-cell formation, an ability that could be encountered the anoxic environment of plant tissue. These results thus provide a lead in understanding where the formation of S-cells in nature is important.

## METHODS

### Strains and media

The wild-type strain *Kitasatospora viridifaciens* DSM40239 ^18^ was obtained from DSMZ. Spores were harvested after 7 days of growth on MYM agar medium ^35^. Vegetative mycelium was grown in the liquid medium TSBS (BD^TM^ Tryptic Soy Broth supplemented with 10% (w/v) sucrose). Liquid-grown cultures were incubated at 30°C, in 100 mL flasks equipped with coils while shaking at 200 rpm. To promote the extrusion of S-cells, high osmotic L-phase broth (LPB) or the solid L-phase medium (LPMA) were used ^17^. All cultures were incubated at 30°C and liquid cultures were grown in 100 mL flasks without coil and gently shaken at 100 rpm.

### Growth of micro-aerobic cultures

Spores or germinated spores of *K. viridifaciens* strains were inoculated on solid LPMA or MYM medium, after which the plates were placed inside a 2.5 L sealed jar (BBL^TM^ GasPak^TM^ from BD Biosciences or from Merck) under micro-aerobic conditions created by using an oxygen-absorbing AnaeroGen sachet (Oxoid). The micro-aerobic atmosphere was monitored using methylene blue indicator strips. The jar was placed in an incubator set at 30°C for two weeks before the strains were further investigated.

### Construction of the Δ*filP* mutant

The *K. viridifaciens* DSM40239 homologue of *filP* (BOQ63_RS19300) was identified by BLAST analysis using the FilP amino acid sequence of *Streptomyces coelicolor* (SCO5396). To create a Δ*filP* mutant, the *filP* gene was replaced with an apramycin-*loxP* cassette (*aac*(3)IV) via the unstable shuttle vector pWHM3-*oriT* ^36,37^ containing the flanking regions of the BOQ63_RS19300 gene (Supplementary Fig. 1A). The flanking regions ranged from −780 bp to +11 upstream and from +916 to +2278 downstream relative to the start codon of the *filP* gene. The PCR-amplified region of the upstream flank of *filP* was cut with restriction enzymes EcoRI and XbaI, whereas the downstream flank of *filP* was cut with XbaI and HindIII, to enable ligation into vector pWHM3-oriT cut with EcoRI and HindIII. The apramycin cassette was inserted in between the *filP* flanks via a XbaI restriction digest and ligation, resulting in vector pWHM3-oriT-*filP*. This vector was verified using a restriction digest with combinations of the abovementioned restriction enzymes (see Supplementary Fig. 1C). The vector was introduced into the *K. viridifaciens* wild-type via conjugation with *E. coli* ET12567/pUZ8002 ^38^, followed by selection for ex-conjugants that were apramycin resistance and thiostrepton sensitive. The successful deletion of the *filP* gene was verified by PCR (Supplementary Fig. 1D). Mutant M1 was chosen and further used in this study.

### Fluorescence and time-lapse microscopy

To stain nucleic acids, SYTO9 (Molecular Probes™) was added to samples obtained from liquid LPB or TSBS broth to a final concentration of 0.5 μM. To fluorescently label lipids, FM5-95 (Molecular Probes™, Invitrogen) or SynapseRed C2M (PromoKine, PromoCell GmbH) was used to a final concentration of 0.02 μg mL^-1^. The fluorescently stained cells were placed in a 35 mm imaging μ-dish (Ibidi^®^) and covered by an agar pad. The samples were analyzed using an inverted Zeiss Axio Observer Z1 confocal microscope equipped with an incubation chamber and Temperature Module S (PECON) stage-top set to 30°C for time-lapse recordings. Fluorescent microscopy images were collected using Zen software (Zeiss) and were viewed, analyzed and edited using ImageJ/FIJI ^39^.

### Cryo-electron microscopy and tomography

*K. viridifaciens* was cultured in liquid LPB medium and after 6 (cryo-ET) or 16 hours (cryo-TEM) 3,5 μl of the culture was mixed with 10 nm gold fiducial markers (Cell Microscopy Core, UMC Utrecht) and applied onto glow-discharged R2/2 200 mesh holey carbon EM grids (Quantifoil). The grid was then rapidly plunged in liquid ethane using the Leica GP automated freeze-plunger, generating a vitrified sample. The vitrified samples were observed using a 626 side-entry cryo-holder (Gatan) and a 120 kV Talos cryo-TEM (Thermo Fisher Scientific) equipped with a Lab6 electron source and Ceta detector. Cryo-TEM tilt series for tomography were collected at the Netherlands Center for Nanoscopy (NeCEN, Leiden) using a 300 kV Titan Krios cryo-TEM (Thermo Fisher Scientific) equipped with a FEG electron gun, GIF energy filter and K2 direct electron detector (Gatan). Tilt series were collected ranging from −60 to +60 with an increment of 2 degrees, and a pixel size of 4.241 A. The data was collected using UCSF software ^40^ and reconstructed to tomograms using IMOD software ^41^.

### Data analysis

Cryo-TEM data was used to determine the cell wall thickness of *K. viridifaciens* hyphae grown under high osmotic stress (LPB medium, 20% sucrose) and to compare with hyphae grown under non-stressed conditions (LPB, 0% sucrose). Of both growth conditions, 15 hyphal tips were imaged. Per tip, three measurements of the cell wall thickness were obtained using ImageJ/FIJI ^39^.

## ACKNOWLEDGEMENTS

We would like to thank Helga van der Heul (Institute of Biology, Leiden University, the Netherlands) for providing the pWHM3-*oriT* vector. The cryo-ET work in this study has been supported by iNEXT, PID:2265, funded by the Horizon2020 programme of the European Commission. Work in the Claessen lab is funded by NWO.

## REFERENCES

1 Höltje, J. V. Growth of the stress-bearing and shape-maintaining murein sacculus of *Escherichia coli*. Microbiol Mol Biol Rev 62, 181–203 (1998).

2 Typas, A., Banzhaf, M., Gross, C. A. & Vollmer, W. From the regulation of peptidoglycan synthesis to bacterial growth and morphology. Nat Rev Microbiol 10, 123–136, doi:10.1038/nrmicro2677 (2012).

3 Pazos, M., Peters, K. & Vollmer, W. Robust peptidoglycan growth by dynamic and variable multi-protein complexes. Curr Opin Microbiol 36, 55–61, doi:10.1016/j.mib.2017.01.006 (2017).

4 van Teeffelen, S. & Renner, L. D. Recent advances in understanding how rod-like bacteria stably maintain their cell shapes. F1000Res 7, 241, doi:10.12688/f1000research.12663.1 (2018).

5 Flärdh, K., Richards, D. M., Hempel, A. M., Howard, M. & Buttner, M. J. Regulation of apical growth and hyphal branching in *Streptomyces*. Curr Opin Microbiol 15, 737–743, doi:10.1016/j.mib.2012.10.012 (2012).

6 Holmes, N. A. et al. Coiled-coil protein Scy is a key component of a multiprotein assembly controlling polarized growth in *Streptomyces*. Proc Natl Acad Sci US A 110, E397–406, doi:10.1073/pnas.1210657110 (2013).

7 Hempel, A. M., Wang, S. B., Letek, M., Gil, J. A. & Flärdh, K. Assemblies of DivIVA mark sites for hyphal branching and can establish new zones of cell wall growth in *Streptomyces coelicolor*. J Bacteriol 190, 7579–7583, doi:10.1128/JB.00839-08 (2008).

8 Fuchino, K. et al. Dynamic gradients of an intermediate filament-like cytoskeleton are recruited by a polarity landmark during apical growth. Proc Natl Acad Sci U S A 110, E1889–1897, doi:10.1073/pnas.1305358110 (2013).

9 Flärdh, K. Essential role of DivIVA in polar growth and morphogenesis in *Streptomyces coelicolor* A3(2). Mol Microbiol 49, 1523–1536 (2003).

10 Bagchi, S., Tomenius, H., Belova, L. M. & Ausmees, N. Intermediate filament-like proteins in bacteria and a cytoskeletal function in *Streptomyces*. Mol Microbiol 70, 1037–1050, doi:10.1111/j.1365-2958.2008.06473.x (2008).

11 Claessen, D. & Errington, J. Cell wall-deficiency as a coping strategy for stress. 7rends in Microbiology (2019).

12 Dörr, T. et al. A cell wall damage response mediated by a sensor kinase/response regulator pair enables beta-lactam tolerance. Proc Natl Acad Sci U S A 113, 404–409, doi:10.1073/pnas.1520333113 (2016).

13 Monahan, L. G. et al. Rapid conversion of *Pseudomonas aeruginosa* to a spherical cell morphotype facilitates tolerance to carbapenems and penicillins but increases susceptibility to antimicrobial peptides. Antimicrob Agents Chemother 58, 1956–1962, doi:10.1128/AAC.01901-13 (2014).

14 Kawai, Y., Mickiewicz, K. & Errington, J. Lysozyme counteracts β-Lactam antibiotics by promoting the emergence of L-form bacteria. Cell 172, 1038–1049.e1010, doi:https://doi.org/10.1016/j.cell.2018.01.021 (2018).

15 Leaver, M., Dominguez-Cuevas, P., Coxhead, J. M., Daniel, R. A. & Errington, J. Life without a wall or division machine in *Bacillus subtilis*. Nature 457, 849–853, doi:10.1038/nature07742 (2009).

16 Allan, E. J., Hoischen, C. & Gumpert, J. Bacterial L-forms. Adv Appl Microbiol 68, 1–39, doi:10.1016/S0065-2164(09)01201-5 (2009).

17 Ramijan, K. et al. Stress-induced formation of cell wall-deficient cells in filamentous actinomycetes. Nat Commun 9, 5164, doi:https://doi.org/10.1101/094037 (2018).

18 Ramijan, K., van Wezel, G. P. & Claessen, D. Genome sequence of the filamentous actinomycete *Kitasatospora viridifaciens*. Genome Announc 5, e01560–01516, doi:10.1128/genomeA.01560-16 (2017).

19 Harold, F. M. Force and compliance: rethinking morphogenesis in walled cells. Fungal Genet Biol 37, 271–282, doi:10.1016/s1087-1845(02)00528-5 (2002).

20 Koch, A. L. & Doyle, R. J. Inside-to-outside growth and turnover of the wall of gram-positive rods. J Theor Biol 117, 137–157, doi:10.1016/s0022-5193(85)80169-7 (1985).

21 Lew, R. R. How does a hypha grow? The biophysics of pressurized growth in fungi. Nat Rev Microbiol 9, 509–518, doi:10.1038/nrmicro2591 (2011).

22 Steinberg, G. Hyphal growth: a tale of motors, lipids, and the Spitzenkörper. Eukaryot Cell 6, 351–360, doi:10.1128/EC.00381-06 (2007).

23 Pilizota, T. & Shaevitz, J. W. Plasmolysis and cell shape depend on solute outer-membrane permeability during hyperosmotic shock in *E. coli*. Biophys J 104, 2733–2742, doi:10.1016/j.bpj.2013.05.011 (2013).

24 Rojas, E. R. & Huang, K. C. Regulation of microbial growth by turgor pressure. Curr Opin Microbiol 42, 62–70, doi:10.1016/j.mib.2017.10.015 (2018).

25 Korber, D. R., Choi, A., Wolfaardt, G. M. & Caldwell, D. E. Bacterial plasmolysis as a physical indicator of viability. Appl Environ Microbiol 62, 3939–3947 (1996).

26 Fuchino, K., Flärdh, K., Dyson, P. & Ausmees, N. Cell-biological studies of osmotic shock response in *Streptomyces* spp. J Bacteriol 199, e00465–00416, doi:10.1128/JB.00465-16 (2017).

27 Bibb, M. J., Molle, V. & Buttner, M. J. σ(BldN), an extracytoplasmic function RNA polymerase sigma factor required for aerial mycelium formation in *Streptomyces coelicolor* A3(2). J Bacteriol 182, 4606–4616 (2000).

28 Kawai, Y. et al. Cell growth of wall-free L-form bacteria is limited by oxidative damage. Curr Biol 25, 1613–1618, doi:10.1016/j.cub.2015.04.031 (2015).

29 Kawai, Y. et al. Crucial role for central carbon metabolism in the bacterial L-form switch and killing by beta-lactam antibiotics. Nat Microbiol, doi:10.1038/s41564-019-0497-3 (2019).

30 van der Meij, A. et al. Inter- and intracellular colonization of *Arabidopsis* roots by endophytic actinobacteria and the impact of plant hormones on their antimicrobial activity. Antonie Van Leeuwenhoek 111, 679–690, doi:10.1007/s10482-018-1014-z (2018).

31 Larsen, M., Santner, J., Oburger, E., Wenzel, W. W. & Glud, R. N. O2 dynamics in the rhizosphere of young rice plants *(Oryza sativa* L.) as studied by planar optodes. Plant Soil 390, 279–292, doi:10.1007/s11104-015-2382-z (2015).

32 Verma, S. K. & White, J. F. Indigenous endophytic seed bacteria promote seedling development and defend against fungal disease in browntop millet *(Urochloa ramosa* L.). JAppl Microbiol 124, 764–778, doi:10.1111/jam.13673 (2018).

33 Verma, S. K. et al. Bacterial endophytes from rice cut grass *(Leersia oryzoides* L.) increase growth, promote root gravitropic response, stimulate root hair formation, and protect rice seedlings from disease. Plant and Soil 422, 223–238, doi:10.1007/s11104-017-3339-1 (2018).

34 Ferguson, C. M. J., Booth, N. A. & Allan, E. J. An ELISA for the detection of *Bacillus subtilis* L-form bacteria confirms their symbiosis in strawberry. Lett Appl Microbiol 31, 390–394 (2000).

35 Stuttard, C. Temperate phages of *Streptomyces venezuelae:* lysogeny and host specificity shown by phages SV1 and SV2. J Gen Microbiol 128, 115–121 (1982).

36 Vara, J., Lewandowska-Skarbek, M., Wang, Y. G., Donadio, S. & Hutchinson, C. R. Cloning of genes governing the deoxysugar portion of the erythromycin biosynthesis pathway in *Saccharopolyspora erythraea (Streptomyces erythreus)*. J Bacteriol 171, 5872–5881 (1989).

37 Światek, M. A. et al. The ROK family regulator Rok7B7 pleiotropically affects xylose utilization, carbon catabolite repression, and antibiotic production in *Streptomyces coelicolor*. J Bacteriol 195, 1236–1248, doi:10.1128/JB.02191-12 (2013).

38 Kieser, T., Bibb, M. J., Buttner, M. J., Chater, K. F. & Hopwood, D. A. Practical Streptomyces genetics. (The John Innes Foundation, 2000).

39 Schindelin, J. et al. Fiji: an open-source platform for biological-image analysis. Nat Methods 9, 676–682, doi:10.1038/nmeth.2019 (2012).

40 Zheng, Q. S., Braunfeld, M. B., Sedat, J. W. & Agard, D. A. An improved strategy for automated electron microscopic tomography. J Struct Biol 147, 91–101, doi:10.1016/j.jsb.2004.02.005 (2004).

41 Kremer, J. R., Mastronarde, D. N. & McIntosh, J. R. Computer visualization of three-dimensional image data using IMOD. J Struct Biol 116, 71–76, doi:10.1006/jsbi.1996.0013 (1996).

