## Supplementary Dataset for "Formation of wall-less cells in *Kitasatospora viridifaciens* requires cytoskeletal protein FilP in oxygen-limiting conditions"

**SUPPLEMENTARY DATA**

**Supplementary Table 1. Quantification of S-cell-related stress response events at the hyphal tip.** Germinated *K. viridifaciens* spores were fluorescently labelled with SYTO9 (nucleic acids) and FM5-95 (lipids) and were grown under high osmotic conditions. The hyphal tips were imaged, examined and categorized. The average percentage per category is visually represented in the graph of Fig. 2.

| **Replica** | **# Tips** | **(A) No extrusion** | **(B) Membrane blebbing** | **(C) Empty vesicles** | **(D) S-cells** | **(E) Empty vesicles and S-cells** |
|---|---|---|---|---|---|---|
| **#1** | 102 | 33 | 14 | 3 | 8 | 42 |
| **#2** | 99 | 35 | 16 | 12 | 19 | 17 |
| **#3** | 84 | 19 | 14 | 20 | 7 | 39 |
| **Average** | | 29 | 15 | 12 | 11 | 33 |
| **SEM** | | 5,0 | 0,66 | 5,1 | 3,9 | 7,8 |

**Supplementary Table 2. Cell wall thickness of *K. viridifaciens* hyphae in the absence and presence of 20% sucrose.** The thickness was measures at three positions per tip.

| Hyphal tip | Condition | Cell wall thickness (nm) #1 | #2 | #3 | Average thickness (nm) |
|---|---|---|---|---|---|
| 1 | 0% sucrose | 41,17 | 39,85 | 39,33 | 40,12 |
| 2 | 0% sucrose | 37,24 | 38,54 | 34,12 | 36,63 |
| 3 | 0% sucrose | 44,42 | 42,75 | 42,40 | 43,19 |
| 4 | 0% sucrose | 45,21 | 44,14 | 44,43 | 44,59 |
| 5 | 0% sucrose | 48,34 | 43,22 | 39,13 | 43,56 |
| 6 | 0% sucrose | 39,20 | 37,88 | 33,92 | 37,00 |
| 7 | 0% sucrose | 49,29 | 46,45 | 48,23 | 47,99 |
| 8 | 0% sucrose | 41,16 | 38,40 | 40,09 | 39,88 |
| 9 | 0% sucrose | 36,27 | 33,89 | 36,55 | 35,57 |
| 10 | 0% sucrose | 46,38 | 40,93 | 46,09 | 44,47 |
| 11 | 0% sucrose | 46,21 | 46,72 | 46,35 | 46,43 |
| 12 | 0% sucrose | 44,62 | 40,77 | 50,07 | 45,15 |
| 13 | 0% sucrose | 39,91 | 41,87 | 43,06 | 41,61 |
| 14 | 0% sucrose | 50,52 | 49,29 | 50,03 | 49,95 |
| 15 | 0% sucrose | 43,46 | 41,94 | 48,62 | 44,67 |
| | **Average** | | | | **42,72** |
| | **SEM** | | | | **1,05** |

| Hyphal tip | Condition | Cell wall thickness (nm) #1 | #2 | #3 | Average thickness (nm) |
|---|---|---|---|---|---|
| 1 | 20% sucrose | 25,68 | 34,48 | 35,95 | 32,04 |
| 2 | 20% sucrose | 26,22 | 29,33 | 26,96 | 27,50 |
| 3 | 20% sucrose | 33,59 | 31,90 | 33,68 | 33,06 |
| 4 | 20% sucrose | 38,10 | 38,23 | 38,66 | 38,33 |
| 5 | 20% sucrose | 38,00 | 35,83 | 35,64 | 36,49 |
| 6 | 20% sucrose | 29,98 | 26,93 | 28,07 | 28,33 |
| 7 | 20% sucrose | 31,16 | 32,88 | 30,90 | 31,65 |
| 8 | 20% sucrose | 27,55 | 19,94 | 19,44 | 22,31 |
| 9 | 20% sucrose | 30,23 | 33,94 | 33,41 | 32,53 |
| 10 | 20% sucrose | 35,68 | 41,33 | 39,60 | 38,87 |
| 11 | 20% sucrose | 27,49 | 26,79 | 34,97 | 29,75 |
| 12 | 20% sucrose | 27,43 | 32,27 | 26,34 | 28,68 |
| 13 | 20% sucrose | 35,45 | 30,63 | 33,26 | 33,11 |
| 14 | 20% sucrose | 25,98 | 25,23 | 27,33 | 26,18 |
| 15 | 20% sucrose | 29,83 | 23,85 | 29,80 | 27,83 |
| | **Average** | | | | **31,11** |
| | **SEM** | | | | **1,14** |

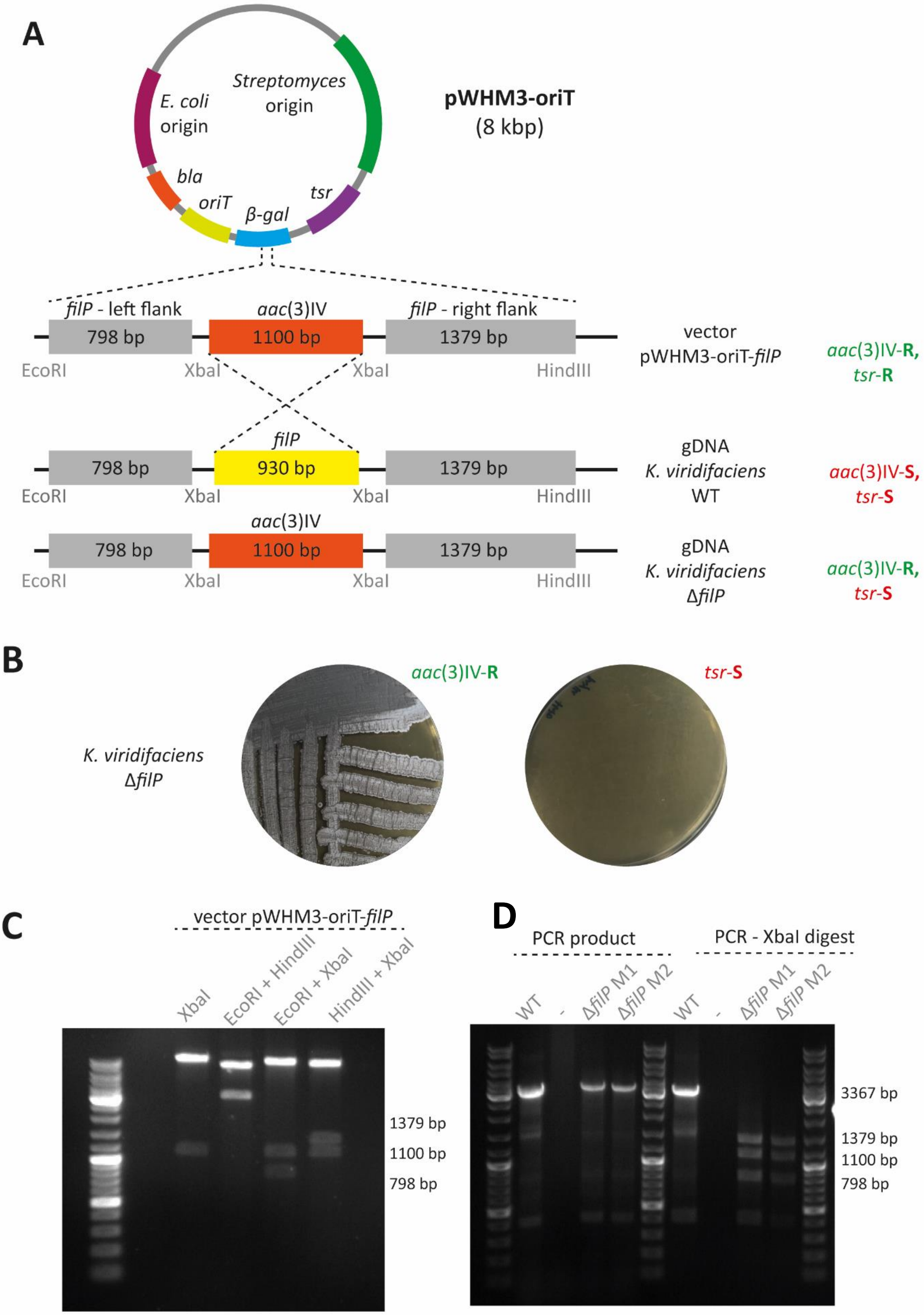


**Supplementary Figure 1. Construction of the Δ*filP* mutant in *K. viridifaciens***

To create a Δ*filP* mutant, the left (upstream) and right (downstream) regions of *filP* were PCR amplified and together with an apramycin resistance cassette (*aac*(3)IV) inserted into pWHM3-*oriT* holding a thiostrepton resistance cassette (*tsr*). Homologous recombination leads to the replacement of the *filP* gene by the *aac*(3)IV apramycin resistance cassette and loss of the pWHM3-*oriT* vector. As a consequence, the Δ*filP* mutant strain is apramycin resistant and thiostrepton sensitive. The pWHM3-*oriT* vector was verified prior to introduction into *K. virdifaciens* (C). (D) PCR amplification of the *filP*-flanking regions and a subsequent digest confirm the successful creation of the Δ*filP* strain.

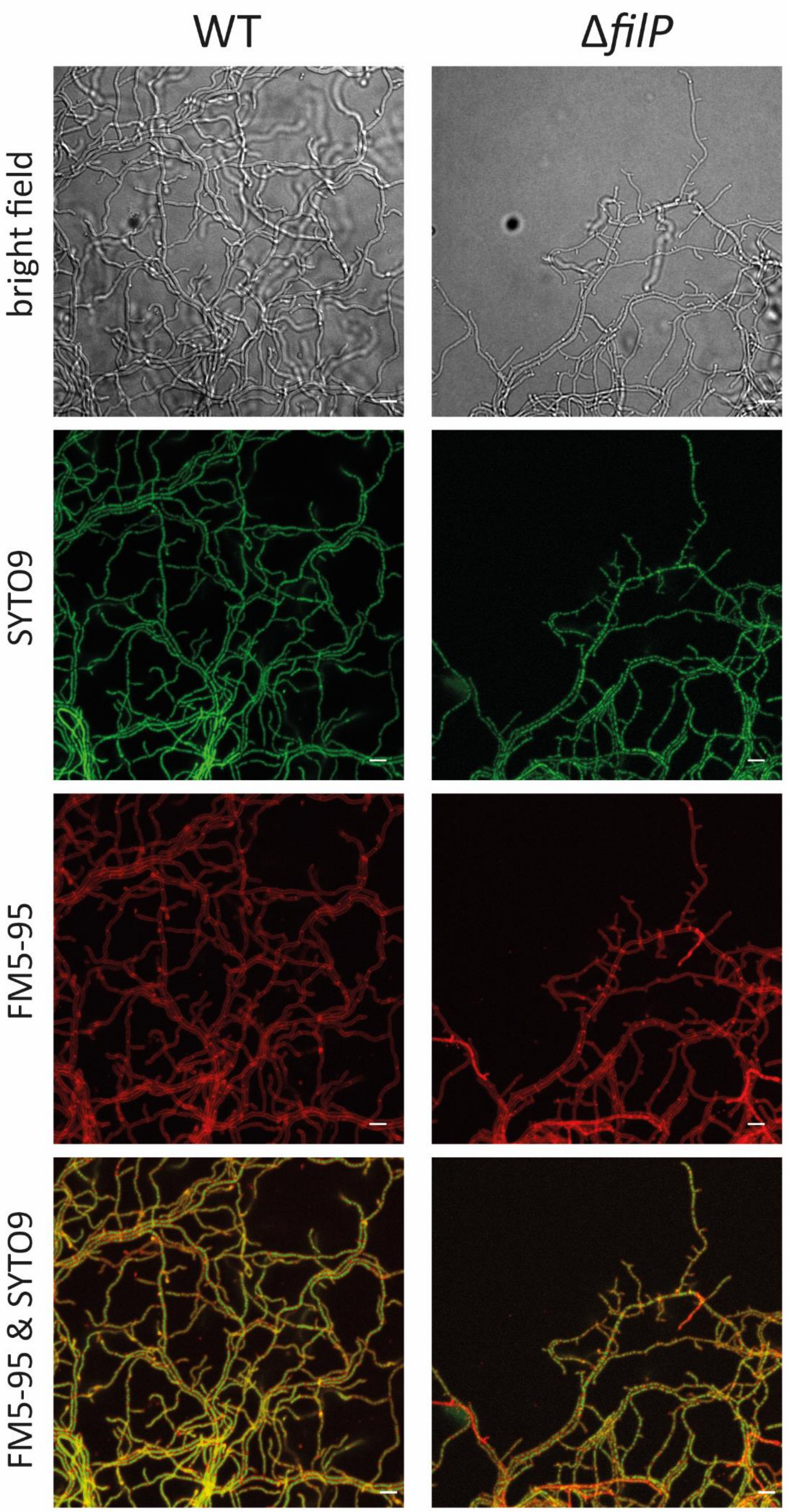


**Supplementary Figure 2. Morphology of the *K. virdifaciens* wild-type and Δ*filP* strains underneath an MYM agar pad**

Germinated *K. viridifaciens* spores were labelled with SYTO9 (nucleic acids, green) and FM5-95 (lipids, red) and grown underneath an MYM agar pad. After 18 hours of growth, the hyphal tips were imaged. Scale bars represent 5 µm.

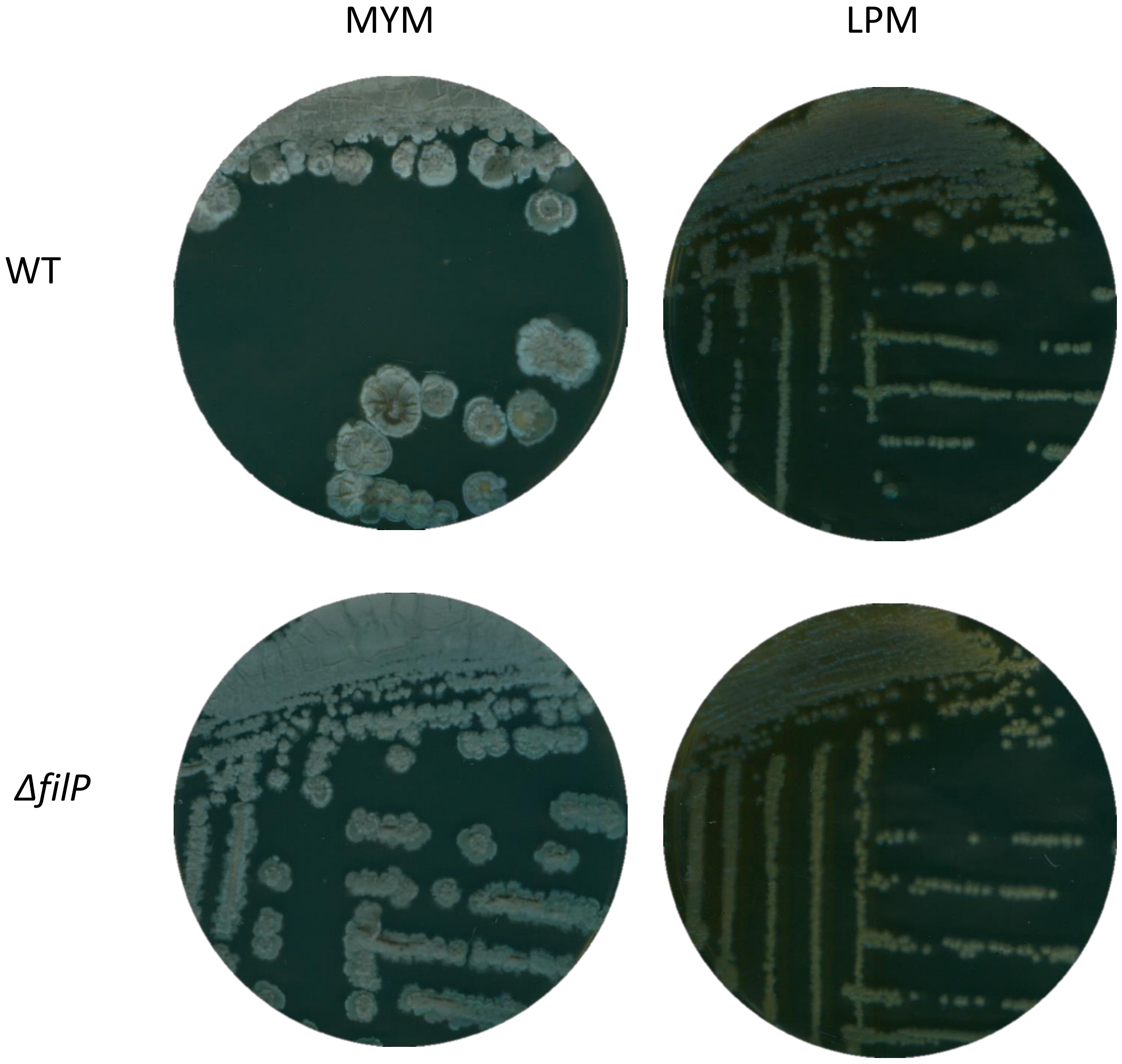


**Supplementary Figure 3. Morphology of the *K. virdifaciens* wild-type and Δ*filP* strains on solid medium in the presence and absence of osmotic stress**

Morphology of the *K. viridifaciens* wild-type and Δ*filP* strains grown on MYM (no osmotic stress) and LPMA (osmotic stress) agar plates for 7 days. Grey-pigmented spores are formed on MYM agar, whereas on LPMA agar both strains produce slimy colonies associated with S-cell formation.

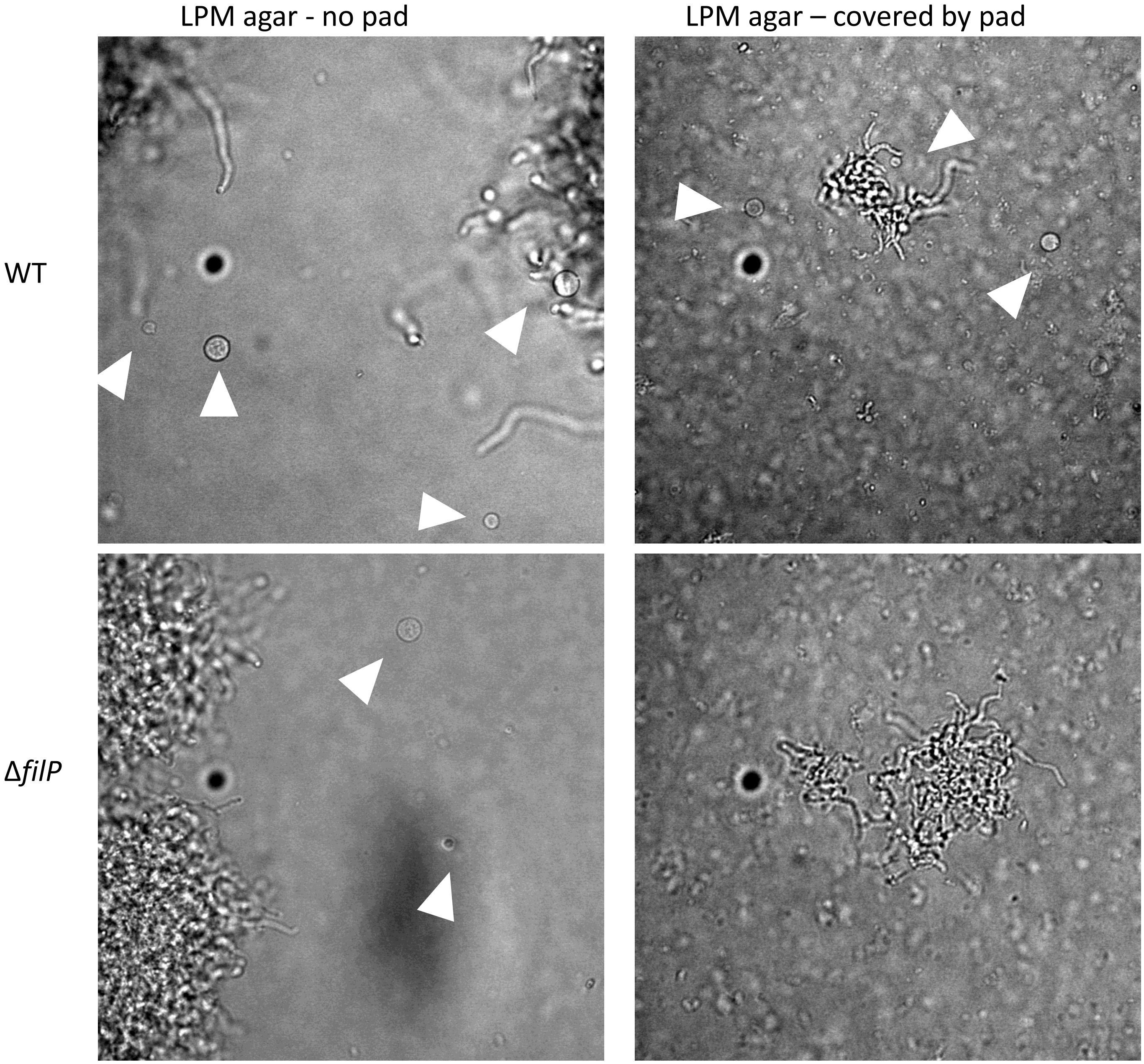


**Supplementary Figure 4. Growth of *K. viridifaciens* germlings covered by agar pads**
Germinated spores of the *K. viridifaciens* wild-type and Δ*filP* strains were cultured on LPMA medium and were either covered by an LPMA agar pad or left to grow without a pad. The wild-type strain forms S-cells in both conditions, while the Δ*filP* strain only forms S-cells when no agar pad is present.

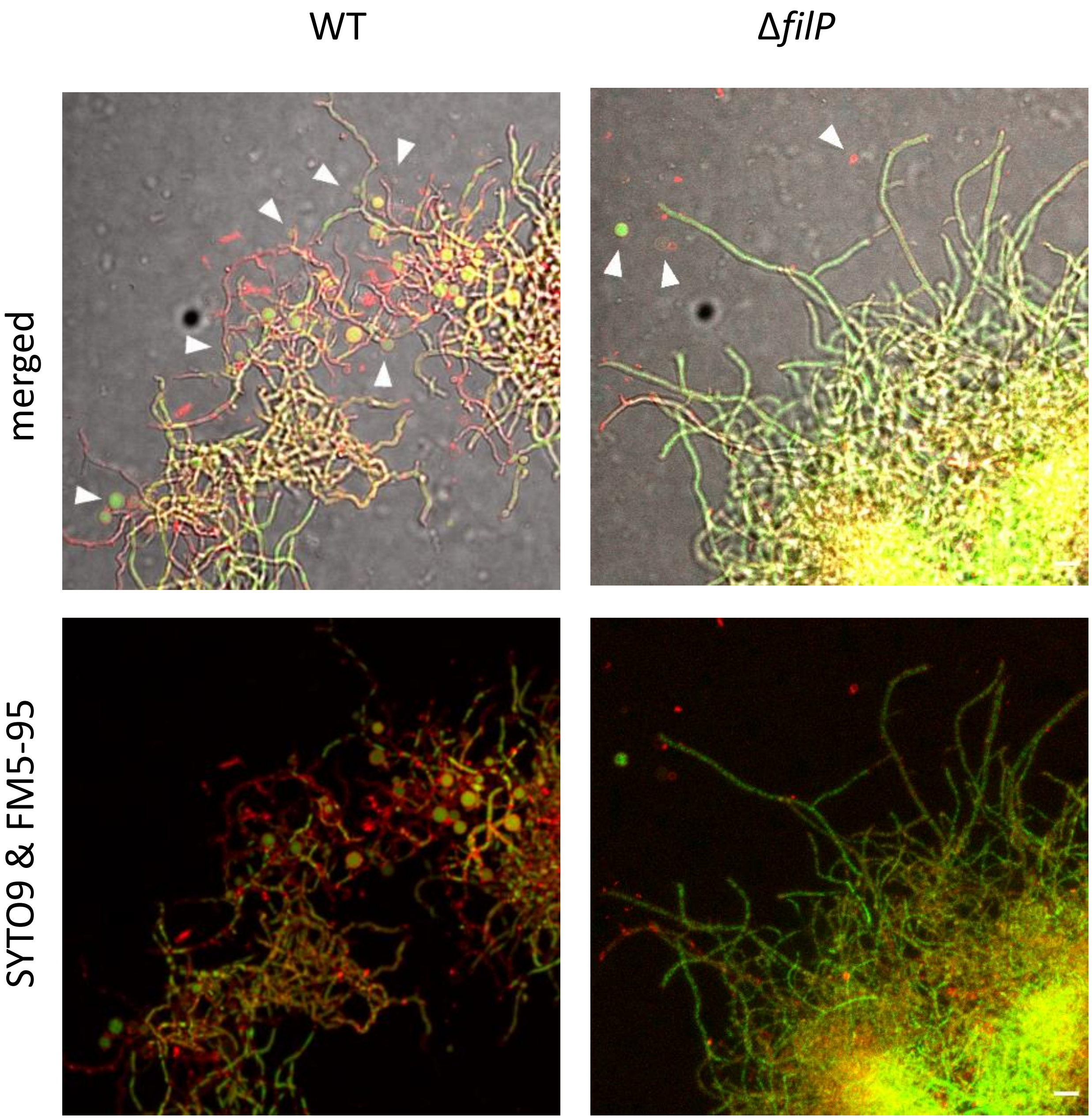


**Supplementary Figure 5. Morphology of the *K. virdifaciens* wild-type and Δ*filP* strains in liquid medium in the presence of osmotic stress**
Germinated *K. viridifaciens* spores were cultured in liquid LPB medium for 18 hours. Upon observation, the cultures were labelled with SYTO9 (nucleic acids, green) and FM5-95 (lipids, red). Arrows highlight the formation of S-cells. Scale bars represent 5 µm.

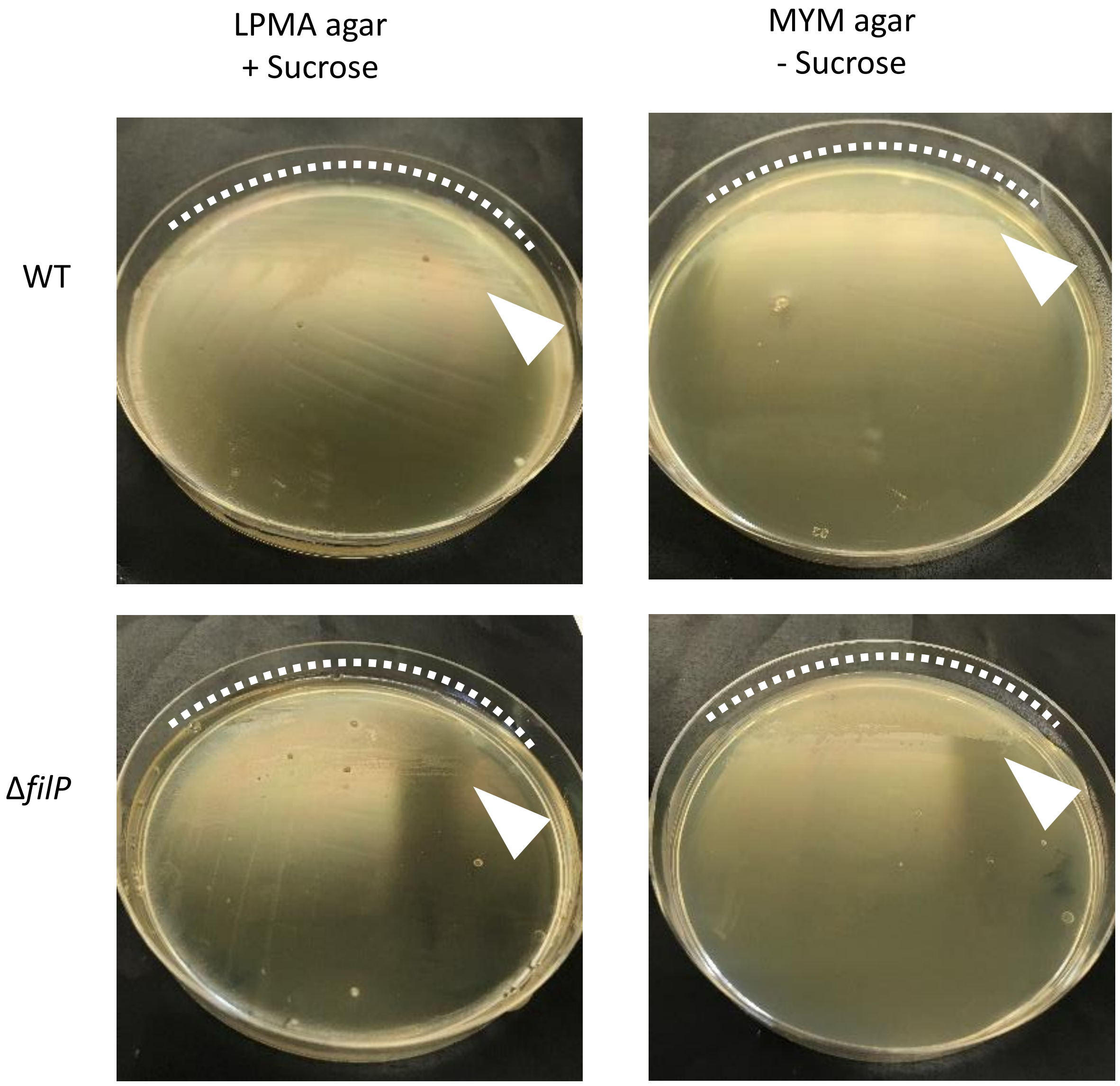


**Supplementary Figure 6. Growth of *K. virdifaciens* in micro-aerobic conditions**
Spores of *K. viridifaciens* strains were inoculated in micro-aerobic conditions on LPMA (osmotic stress) and MYM (no osmotic stress) medium. After two weeks, growth was observed on all plates, indicated by the dashed lines and arrowheads.

**SUPPLEMENTARY MOVIE LEGENDS**

**Supplementary Movie 1A / 1B**

Time-lapse recording of germinated *K. viridifaciens* spores stained with STYO9 (nucleic acid, green) and FM5-95 (lipids, red) and grown under high osmotic conditions using a LPMA agar pad. Movies 1A and 1B represent two different areas of the recording. The 'S' indicates the spore, the arrow heads highlight the blebbing of membrane, the asterisks indicates S-cell formation. Recording time is represented in hours:minutes, scale bar is 5 µm.

**Supplementary Movie 2**

Time-lapse recording of germinated *K. viridifaciens* WT spores grown under non-stressed conditions, using a MYM agar pad. Recording time is represented in hours:minutes, scale bar is 5 µm.

**Supplementary Movie 3**

Time-lapse recording of germinated *K. viridifaciens* Δ*filP* spores grown under non-stressed conditions, using a MYM agar pad. Recording time is represented in hours:minutes, scale bar is 5 µm.

**Supplementary Movie 4**

Time-lapse recording of germinated *K. viridifaciens* wild-type spores grown under high osmotic conditions, using a LPMA agar pad. Recording time is represented in hours:minutes, scale bar is 5 µm.

**Supplementary Movie 5**

Time-lapse recording of germinated *K. viridifaciens* Δ*filP* spores grown under high osmotic conditions, using a LPMA agar pad. Recording time is represented in hours:minutes, scale bar is 5 µm.
